# Tissue-guided LASSO for prediction of clinical drug response using preclinical samples

**DOI:** 10.1101/724310

**Authors:** Edward W Huang, Ameya Bhope, Jing Lim, Saurabh Sinha, Amin Emad

## Abstract

Prediction of clinical drug response (CDR) of cancer patients, based on their clinical and molecular profiles obtained prior to administration of the drug, can play a significant role in individualized medicine. Machine learning models have the potential to address this issue, but training them requires data from a large number of patients treated with each drug, limiting their feasibility. While large databases of drug response and molecular profiles of preclinical *in-vitro* cancer cell lines (CCLs) exist for many drugs, it is unclear whether preclinical samples can be used to predict CDR of real patients.

We designed a systematic approach to evaluate how well different algorithms, trained on gene expression and drug response of CCLs, can predict CDR of patients. Using data from two large databases, we evaluated various linear and non-linear algorithms, some of which utilized information on gene interactions. Then, we developed a new algorithm called TG-LASSO that explicitly integrates information on samples’ tissue of origin with gene expression profiles to improve prediction performance. Our results showed that regularized regression methods provide significantly accurate prediction. However, including the network information or common methods of including information on the tissue of origin did not improve the results. On the other hand, TG-LASSO improved the predictions and distinguished resistant and sensitive patients for 7 out of 13 drugs. Additionally, TG-LASSO identified genes associated with the drug response, including known targets and pathways involved in the drugs’ mechanism of action. Moreover, genes identified by TG-LASSO for multiple drugs in a tissue were associated with patient survival. In summary, our analysis suggests that preclinical samples can be used to predict CDR of patients and identify biomarkers of drug sensitivity and survival.

**AUTHOR SUMMARY:** Cancer is among the leading causes of death globally and perdition of the drug response of patients to different treatments based on their clinical and molecular profiles can enable individualized cancer medicine. Machine learning algorithms have the potential to play a significant role in this task; but, these algorithms are designed based the premise that a large number of labeled training samples are available, and these samples are accurate representation of the profiles of real tumors. However, due to ethical and technical reasons, it is not possible to screen humans for many drugs, significantly limiting the size of training data. To overcome this data scarcity problem, machine learning models can be trained using large databases of preclinical samples (e.g. cancer cell line cultures). However, due to the major differences between preclinical samples and real tumors, it is unclear how accurately such preclinical-to-clinical computational models can predict the clinical drug response of cancer patients.

Here, first we systematically evaluate a variety of different linear and nonlinear machine learning algorithms for this particular task using two large databases of preclinical (GDSC) and tumor samples (TCGA). Then, we present a novel method called TG-LASSO that utilizes a new approach for explicitly incorporating the tissue of origin of samples in the prediction task. Our results show that TG-LASSO outperforms all other algorithms and can accurately distinguish resistant and sensitive patients for the majority of the tested drugs. Follow-up analysis reveal that this method can also identify biomarkers of drug sensitivity in each cancer type.

## INTRODUCTION

Cancer is one of the leading causes of death globally and is expected to be the most important obstacle in increasing the life expectancy in the 21^st^ century [1]. Individualized cancer medicine has the potential to revolutionize patient prognosis; however, two major challenges in this area include the prediction of the individual responses to different treatments and the identification of molecular biomarkers of drug sensitivity. While factors such as cancer type or its symptoms have been traditionally used to identify the treatment [2], the development of high throughput sequencing technologies [3] and sophisticated machine learning (ML) approaches present the possibility of individualizing treatment based on molecular ‘omics’ profiles of patients’ tumors [4]. However, due to the technical and ethical challenges of screening individuals against many drugs [5], such models are either trained for only a handful of drugs [6] or are trained using preclinical samples such as 2D cancer cell line cultures (CCLs) [7–10]. In spite of the success of these methods in predicting the drug response of left-out *preclinical* samples using models trained on *preclinical* samples, they have had limited success in predicting the CDR of real patients [9, 11], with some exceptions [12–14].

Various preclinical models of cancer have been developed to enable the study of cancer and its treatment in the laboratory. CCLs, which are 2D cell cultures developed from tumor samples, are one of the least expensive and most studied of these models. Recently, several large-scale studies have cataloged the molecular profiles of thousands of CCLs and their response to hundreds of drugs [15–17]. Although various computational models have been developed to predict the CCLs’ drug response using their molecular profiles [7–9], these models have shown limited success in predicting CDR in real patients. In spite of sporadic successes for a handful of drugs [12, 13], the current belief remains that developing an accurate computational ‘preclinical-to-clinical’ model is extremely difficult if not impossible [5]. Our goal in this study was to perform an unbiased systematic evaluation on a panel of drugs to determine 1) whether regression models trained on *in vitro* preclinical samples can accurately predict the CDR of real patients for each drug and 2) what type of side information (e.g. interaction of the genes, the tissue of origin of samples) might improve the CDR prediction.

To this end, we first formed a computational framework to systematically evaluate the prediction accuracy of different computational methods. We obtained preclinical training samples from the Genomics of Drug Sensitivity in Cancer (GDSC) database [16] and obtained molecular profiles of tumor samples from The Cancer Genome Atlas (TCGA) [18]. We focused on drugs that were shared between these two datasets and utilized the gene expression profiles of samples to predict the drug response, since previous studies have demonstrated gene expression to be most informative for this task [7]. Our analysis showed that regularized linear regression models provide the best performance among various algorithms. In addition, we included prior information on the relationship among genes (in the form of gene interaction networks) using several algorithms; however, this prior information did not improve the prediction.

Next, we developed a novel approach called Tissue-Guided LASSO (TG-LASSO) to explicitly include information on the tissue of origin of samples in the regularized regression model. This method outperformed all other approaches evaluated. Using this method, we showed that the CDR of cancer patients can be accurately predicted using preclinical CCL training samples, for the majority of drugs. More specifically, out of 12 drugs, TG-LASSO accurately separated resistant patients from sensitive patients for 7 drugs. In addition, for each tissue type and drug, TG-LASSO identified a small set of genes that may be used as tissue-specific biomarkers of drug response for each drug. We showed that genes selected by TG-LASSO for prediction of drug response are informative of patient survival when used as a gene signature, and also provide pathway-level insights into mechanisms of drug action. These results emphasize the clinical relevance of molecular profiles of preclinical samples cataloged in large-scale databases and demonstrate the importance of properly including information on the lineage of samples in follow-up analyses.

## RESULTS

### Prediction of clinical drug response of cancer patients using *in vitro* experiments on preclinical cancer cell lines

In this study, our first goal was to determine whether commonly used machine learning algorithms are capable of predicting the clinical drug response (CDR) in cancer patients using computational models trained only on cancer cell lines’ (CCLs) basal gene expression profiles (i.e. before administration of the drug) and their drug response. For this purpose, we identified 23 drugs (Supplementary Table S1) that were administered to patients of The Cancer Genome Atlas (TCGA) [18] and were also present in the Genomics of Drug Sensitivity in Cancer (GDSC) [16] database. We obtained the gene expression profiles of 531 primary tumor samples of TCGA patients (17 different cancer types) who were administered any of these drugs from the Genomic Data Commons [19] (see Methods and Supplementary Table S1). We obtained the carefully collected and curated information on clinical drug response (CDR) of these patients from [6]. Similarly, we obtained the gene expression profiles and the half-maximal inhibitory concentration (IC50) of 979 cancer cell lines (of 55 different tissues) from GDSC (see Supplementary Table S1 for the number of cell lines from each tissue).

We formed a computational framework to systematically evaluate the prediction capability of different algorithms (Fig. 1). In this framework, we first normalize the data and remove batch effects to ensure that the gene expression profiles from these two datasets are comparable (Methods). This is particularly important since GDSC contains microarray gene expression values, while TCGA contains RNA-seq data. We used ComBat [20] for batch effect removal, which has been previously used to successfully remove the batch effect between RNA-seq and microarray data [21] (see Supplementary Fig. S1 for the distribution of samples before and after batch effect removal). Next, we trained a regression model to relate the gene expression profiles of CCLs to their IC50 values for a specific drug. Given this model, we then estimated IC50 values for different patient tumors using their gene expression profiles. Finally, we compared the estimated IC50 values to the true CDR of the tumors of patients treated with the same drug to determine the accuracy of prediction.

**Figure 1:**
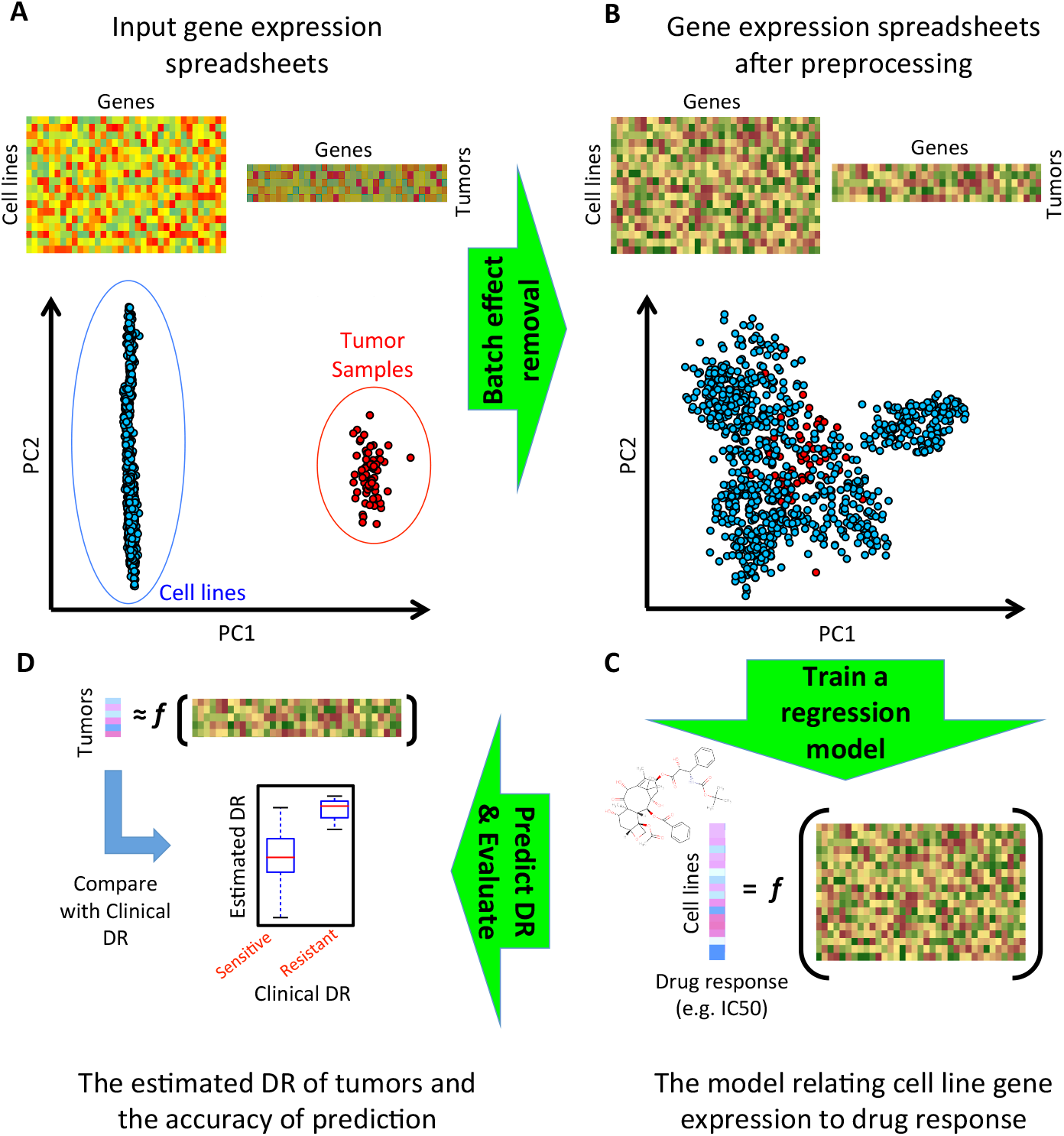
The pipeline used for prediction of clinical drug response of cancer patients using computational models trained on gene expression and drug response of preclinical cell line samples.

We used a one-sided nonparametric Mann Whitney U test to determine whether the estimated IC50 values of resistant tumors (those with CDR of ‘clinical progressive disease’ or ‘stable disease’) are significantly larger than sensitive tumors (those with CDR of ‘partial response’ or ‘complete response’). In this evaluation, we only used 12 drugs that had at least 2 tumor samples in each category of resistant or sensitive and had at least 8 total samples with known CDR. Table 1 shows a summary of the performance of different methods. In this table, we used the combined p-value of all 12 drugs (using Fisher’s method to combine p-values) as a measure to summarize the results of different methods. Table 2 and Supplementary Table S2 contain the detailed performance of LASSO and all other methods, respectively, for prediction of the CDR of each drug. We focused on these methods as they have been previously used for this task (but for fewer drugs and using other datasets), with different degrees of success [12, 22, 23]. Recently, [5] reported a computational model based on ridge regression to predict the CDR of TCGA patients using GDSC training samples. Table 1 also includes the performance of this method using our evaluation, based on the predictions reported in the original paper.

**Table 1:**
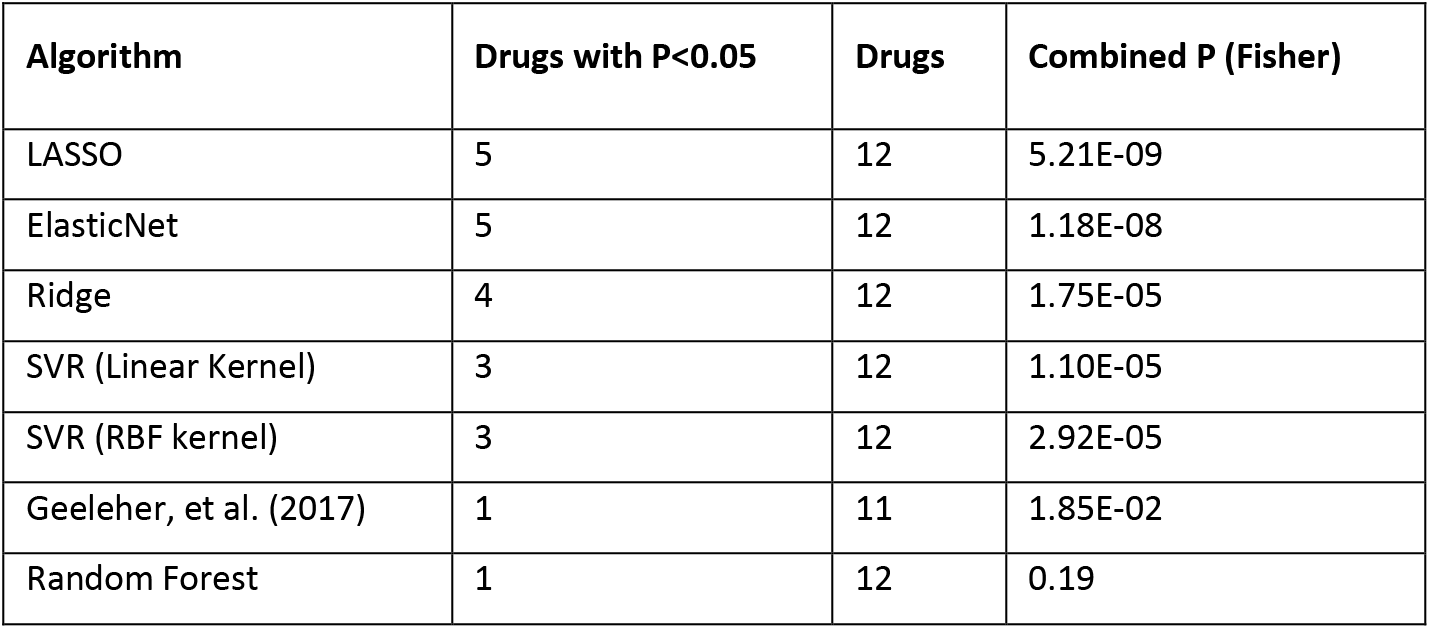
The performance of different algorithms in predicting the CDR of patients using models trained on preclinical CCL samples. The second column shows the number of drugs for which a statistically significant discrimination between resistant and sensitive patients was obtained (one-sided Mann Whitney U test). The third column shows the total number of drugs included in the evaluation, and the fourth column shows the combined p-value (using Fisher’s method) for all the drugs in the analysis.

| Algorithm | Drugs with $P < 0.05$ | Drugs | Combined P (Fisher) |
| --- | --- | --- | --- |
| LASSO | 5 | 12 | 5.21E-09 |
| ElasticNet | 5 | 12 | 1.18E-08 |
| Ridge | 4 | 12 | 1.75E-05 |
| SVR (Linear Kernel) | 3 | 12 | 1.10E-05 |
| SVR (RBF kernel) | 3 | 12 | 2.92E-05 |
| Geeleher, et al. (2017) | 1 | 11 | 1.85E-02 |
| Random Forest | 1 | 12 | 0.19 |

**Table 2:** The performance of LASSO algorithm in predicting the CDR of patients using models trained on preclinical CCL samples. The second column shows the p-value (one-sided Mann Whitney U test) for the predicted IC50 values of sensitive and resistant tumors. The third and fourth columns show the number of resistant and sensitive tumors used in the statistical test.

| Drug | P-value<br>(one-sided) | Num Resistant<br>(PD or SD) | Num Sensitive<br>(CR or PR) |
| --- | --- | --- | --- |
| bicalutamide | 0.34 | 3 | 14 |
| bleomycin | 0.10 | 4 | 46 |
| cisplatin | 6.67E-05 | 25 | 111 |
| docetaxel | 0.98 | 17 | 55 |
| doxorubicin | 3.42E-03 | 7 | 54 |
| etoposide | 7.57E-04 | 10 | 71 |
| gemcitabine | 0.14 | 43 | 37 |
| paclitaxel | 0.62 | 28 | 74 |
| sorafenib | 0.19 | 13 | 2 |
| tamoxifen | 8.82E-03 | 4 | 14 |
| temozolomide | 9.08E-02 | 84 | 11 |
| vinorelbine | 2.10E-03 | 6 | 23 |

These results suggest several important points. First, consistent with the reports in [12, 22], we observed that regularized linear models resulted in the best performance, with LASSO performing the best. Second, we observed that although the method proposed in [5] is based on ridge regression, its performance is inferior to the ridge regression utilized in our study. This is likely due to the difference between the preprocessing and batch effect removal approaches used in the two studies. More specifically, instead of using ComBat to homogenize the gene expression data in the preclinical and clinical samples (as was done in our study and also in [12]), they simply standardized the mean of each gene to zero and its variance to one. This point emphasizes the importance of data preprocessing in pharmacogenomics studies. Third, we observed that for some drugs, the CDR could be accurately predicted independent of the method, while for others, the choice of the method is important. For example, the CDR of cisplatin could be accurately predicted (p<0.05) using six out of the seven methods above (as an example Table 3 shows that 92% of resistant patients are correctly designated using LASSO, while keeping precision at ^~^30% and specificity at ^~^50%). On the other hand, only support vector regression (SVR) with RBF kernel could accurately predict the CDR of temozolomide, suggesting a nonlinear relationship between the gene expression values and the drug response. As another example, the majority of the methods could not predict the CDR of taxane-based chemotherapy agents (docetaxel and paclitaxel). We suspect that this lack of success is due to the existence of various parameters that influence their response, such as tissue dependence or microenvironmental factors [24, 25], which may not be captured using these simple methods trained on gene expression profiles of CCLs. In fact, we later show that including the tissue of origin *explicitly* in the predicting model using TG-LASSO can significantly improve the drug response prediction for paclitaxel.

**Table 3:** The contingency table of predictions using LASSO for cisplatin. The predicted IC50 values were labeled as resistant or sensitive based on the threshold that obtained the highest oddsratio (oddsratio = 10.5, p<0.001).

|  | Predicted Resistant | Predicted Sensitive | Total |
| --- | --- | --- | --- |
| True Resistant | 23 | 2 | 25 |
| True Sensitive | 58 | 53 | 111 |
| Total | 81 | 55 | 136 |

### Including information on gene interactions does not improve CDR prediction

Various studies have suggested that including information on the interaction of the genes (and their protein products) or their involvement in different pathways can improve the accuracy of different bioinformatics tasks [26] such as gene prioritization [27], gene function prediction [28], gene set characterization [29], and tumor subtyping [30]. Since the genes (and their protein products) involved in a drugs mechanism of action biochemically and functionally interact with each other, we sought to determine whether including these interactions could improve CDR prediction. Since linear models provided the best performance in our preliminary analyses (Table 1), we focused on methods that incorporate gene interaction networks into linear predictive models. These included Generalized Elastic Net (GELnet) [31], Network-Induced Classification Kernels (NICK) [32], Sparse Group LASSO (SGL) [33], as well as a method based on LASSO combined with single sample gene set enrichment analysis (ssGSEA) [34] (see Methods). In all cases, we used four gene interaction networks: an experimentally verified network of protein-protein and genetic interactions, a gene co-expression network, and a network built based on text mining from the STRING database [35], as well as the HumanNet integrated network [36] (see Methods and Supplementary Table S1 for details). Table 4 summarizes the results and Supplementary Table S3 provides the details of the evaluations. These results suggest that in this application, incorporating network information using these methods does not improve the prediction compared to linear models (e.g. LASSO) that do not incorporate such information (Table 1). This was in spite of the fact that some of these network-guided methods (e.g. NICK with STRING Text Mining) do improve the performance of within-dataset cross-validation (using only GDSC samples) compared to LASSO (see Supplementary Methods).

**Table 4:** The performance of network-based algorithms in predicting the CDR of patients using models trained on preclinical CCL samples. The third column shows the number of drugs for which a statistically significant discrimination between resistant and sensitive patients was obtained (one-sided Mann Whitney U test). The fourth column shows the total number of drugs included in the evaluation, and the fifth column shows the combined p-value (using Fisher’s method) for all the drugs in the analysis. As a point of comparison, LASSO without the use of any network yielded p-value < 0.05 for five of 12 drugs, with combined p-value of 5.21E-09 (Table 1).

| Network | Algorithm | Drugs with P<0.05 | Drugs | Combined P (Fisher) |
| --- | --- | --- | --- | --- |
| STRING PPI | NICK | 5 | 12 | 1.79E-06 |
|  | GELnet | 5 | 12 | 1.21E-04 |
|  | SGL | 3 | 12 | 1.75E-05 |
|  | ssGSEA-LASSO | 3 | 12 | 6.94E-06 |
| STRING Co-Expression | NICK | 5 | 12 | 2.40E-06 |
|  | GELnet | 5 | 12 | 1.07E-04 |
|  | SGL | 2 | 12 | 2.62E-04 |
|  | ssGSEA-LASSO | 4 | 12 | 1.40E-2 |
| STRING Text Mining | NICK | 5 | 12 | 2.14E-06 |
|  | GELnet | 5 | 12 | 1.23E-04 |
|  | SGL | 3 | 12 | 7.95E-05 |
|  | ssGSEA-LASSO | 3 | 12 | 2.57E-02 |
| HumanNet Integrated Network | NICK | 5 | 12 | 1.09E-06 |
|  | GELnet | 5 | 12 | 1.21E-04 |
|  | SGL | 3 | 12 | 7.69E-04 |
|  | ssGSEA-LASSO | 4 | 12 | 6.06E-07 |

### Incorporating the tissue of origin to improve CDR prediction

Up to this point, we only used the tissue of origin of the preclinical and clinical samples *implicitly* (through their gene expression profiles) by training a single model for a drug on all CCLs of different lineages, and then using this global model to predict the response of patients with different cancer types. However, due to the importance of the tissue of origin in the efficacy of anticancer drugs observed in various studies [37, 38], we sought to determine whether *explicitly* including the tissue of origin would improve the prediction of CDR, and if so, the best method for this inclusion. For our analysis, we focused on variations of LASSO (without including gene interactions), which previously yielded the best performance among all the tested algorithms (Table 1). We matched the lineage of the CCLs with those of cancer patients, identifying 13 shared tissue types.

One of the most common methods of including the tissue of origin in regression analysis is introducing new binary features to each sample, representing whether the sample belongs to that tissue (‘1’) or not (‘0’) [15]. We included 13 such binary features in the analysis (‘method 1’). However, the prediction results of this approach were almost identical to the results of LASSO when not including any tissue information. This is not surprising, since in this application the number of one type of features (i.e. genes) is much larger than the number of the other type of features (i.e. tissue types). As a result, the predicted drug response values will be highly biased by the influence of gene expression data and the tissue of origin’s influence will be overlooked. As an alternative, we trained different LASSO models for each tissue type by restricting the training (CCL) and test (tumor) samples to those originating from the same tissue of interest (‘method 2’). For tumor samples without CCLs with matching tissue, we used all CCLs to train the model. This method resulted in poor performance, with only one drug having a significant p-value and a combined p-value (Fisher’s method) of 0.16. The reason behind this poor performance is the small number of samples in training each model: due to the tissu-especificity condition imposed above, only a small fraction of the total samples are used in training each model, which results in poor generalizability of the models.

To overcome these issues, while explicitly incorporating information on the samples’ tissue of origin, we devised a new approach called <u>T</u>issue-<u>G</u>uided <u>LASSO</u> (TG-LASSO). The idea behind this approach is to use *all* CCLs originating from different tissue types in training the LASSO model, but choose the hyperparameter of the LASSO model, *α*, in a tissue-specific manner. This avoids the issues caused by the small number of training samples in Method 2, while adding a tissue-specific aspect to the training of the model. Since *α* controls the number of features (i.e. genes) used by the LASSO model, this approach allows us to optimally select the number of predictive genes for each tissue type (see Methods for details), yet use all CCLs to train these tissue-specific regression models. This approach resulted in the best performance among all the methods tested, with 7 (out of 12) drugs showing significant discrimination between resistant and sensitive tumors (p<0.05) and a combined p-value (Fisher’s method for all 12 drugs) of 2.25E-10 (Fig. 2, Table 5 and Supplementary Table S4). These results not only show that including the tissue of origin can improve CDR prediction using preclinical samples, but also suggest that the method of utilizing this information has a significant influence on the performance.

**Figure 2:**
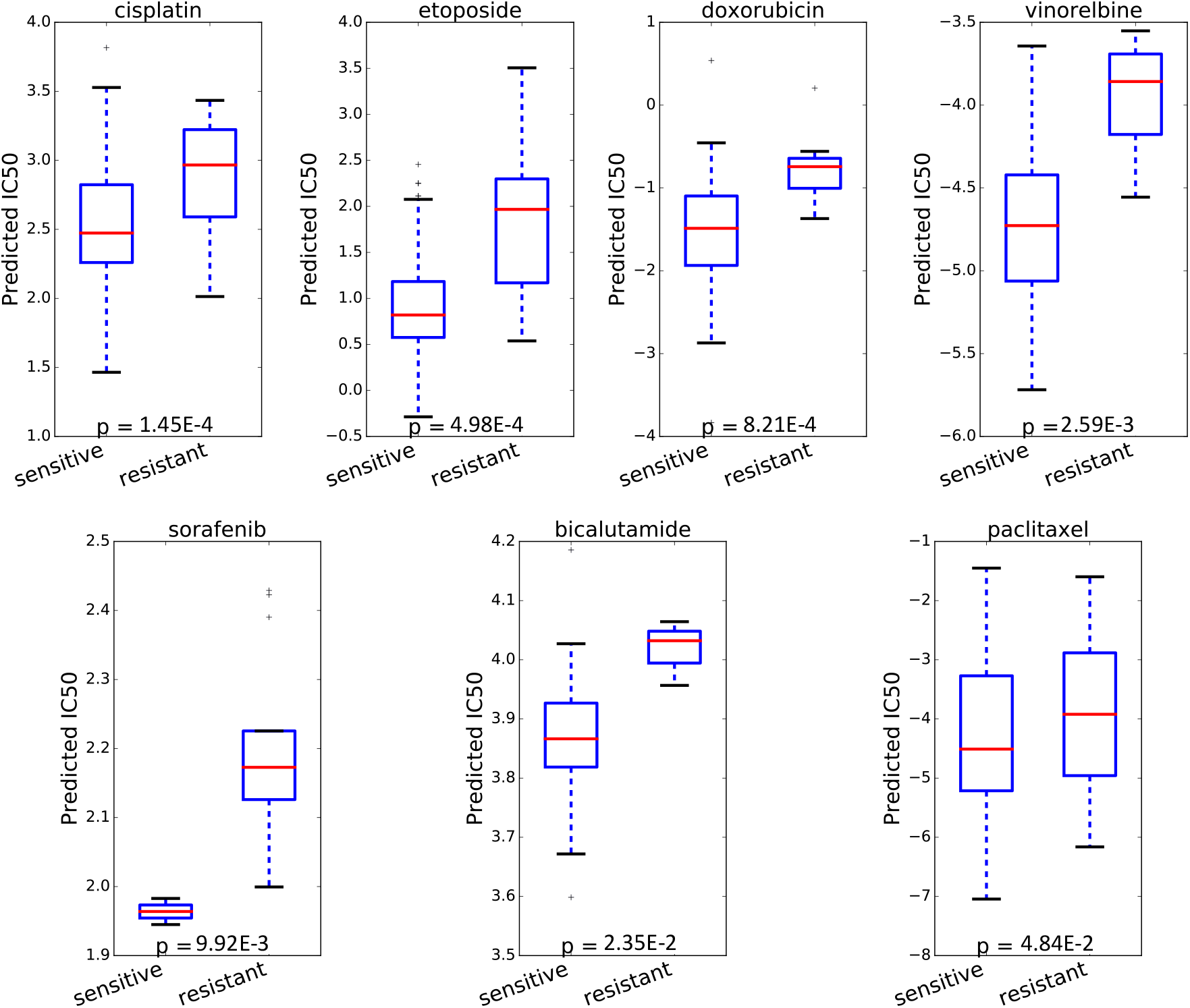
The drug response prediction performance of seven drugs for which TG-LASSO predictions accurately separated sensitive patients from resistant. The box plots reflect the distribution of estimated IC50 values using TG-LASSO for each group of resistant or sensitive patients.

**Table 5:**
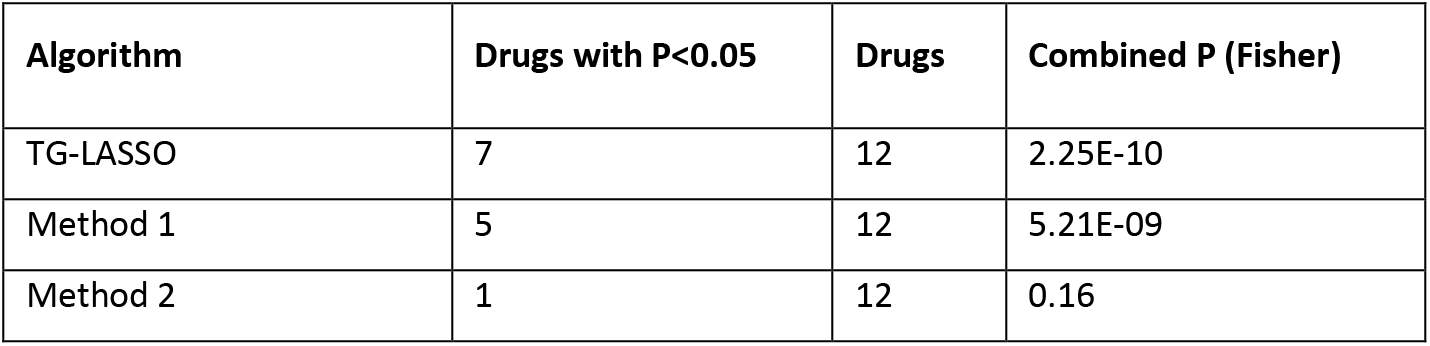
The prediction performance of different approaches that incorporate information on tissue of origin in LASSO. The second column shows the number of drugs for which a statistically significant discrimination between resistant and sensitive patients was obtained (one-sided Mann Whitney U test). The third column shows the total number of drugs included in the evaluation, and the fourth column shows the combined p-value (using Fisher’s method) for all the drugs in the analysis.

| Algorithm | Drugs with P<0.05 | Drugs | Combined P (Fisher) |
| --- | --- | --- | --- |
| TG-LASSO | 7 | 12 | 2.25E-10 |
| Method 1 | 5 | 12 | 5.21E-09 |
| Method 2 | 1 | 12 | 0.16 |

One interesting observation was that paclitaxel, the response of which could not be predicted accurately with the majority of methods reported in Table 1, showed a significant improvement in the response prediction with TG-LASSO (p = 0.048, one-sided Mann Whitney U test), suggesting a prominent role for the tissue of origin in its drug response. On the other hand, the CDR prediction of docetaxel did not improve (p = 0.99), even though docetaxel is also a taxane, like paclitaxel, and these two drugs have a statistically significant correlation in their CCL responses (Spearman rank correlation = 0.38, p = 1.7E-13). We suspected that this difference between the performance of TG-LASSO for docetaxel and paclitaxel is related to how well the CCL panel used for training represents the tumor samples of patients to whom these drugs were administered. To evaluate this, we calculated the similarity between the gene expression profiles of tumor samples to those of CCLs from the same tissue of origin for these drugs. This analysis showed a lower similarity between the docetaxel-administered tumors and CCLs (average cosine similarity = 0.07) compared to paclitaxel-administered tumors and CCLs (average cosine similarity = 0.11). These results provide evidence in favor of our hypothesis that the difference in the performance of TG-LASSO is related to how well the CCLs represent the profile of tumors to which these two drugs were administered.

Since some of the drugs used in our study were administered in combination with other drugs, we asked how well TG-LASSO predicts the CDR in such cases of treatment with drug combinations. For this purpose, we evaluated its CDR prediction for a drug only on patients for whom that drug was administered over a period overlapping their treatment with at least one other drug. We limited our analysis to 9 drugs with at least two samples (patients) in each group (sensitive and resistant) and with at least 8 samples in total. Supplementary Table S5 shows that, consistent with our previous results, TG-LASSO outperforms all other methods, capable of accurately predicting the CDR of 6 (out of 9) drugs (p<0.05, one-sided Mann Whitney U test).

### Characterization of genes identified by TG-LASSO

During its training phase, TG-LASSO automatically selects a subset of genes to be used in the regression model by tuning the hyperparameter *α* introduced above. The number of genes selected in this manner depends on the drug and tissue type for which the model is trained to make response predictions and was found to range between 9 and 808 genes with a median of 174 genes. The genes identified by TG-LASSO included many direct targets of each drug. (For these analyses we used all 23 drugs shared between TCGA and GDSC and not just those with a large number of samples in TCGA). For example, EGFR, which is a direct target of both cetuximab and gefitinib [39], was selected by this algorithm when trained to predict response of these drugs in each of the 13 tissue types (Supplementary Table S6). Similarly, FLT3, a target of the drugs sorafenib and sunitinib [39], was selected by TG-LASSO for predicting response to these drugs in 13 and 12 tissues, respectively. In addition to direct targets, many of the identified genes have been shown to be indirect targets of these drugs and to be involved in their mechanism of action. For example DNER, a gene identified by TG-LASSO for all tissue types, has been shown to be significantly upregulated in response to this drug in NCI-H526 cell lines [40].

More importantly, the knockdown or overexpression of many of the identified genes has been shown to influence the sensitivity of cancer cells to these drugs. For example, the shRNA knockdown of CHI3L1, a gene identified for etoposide and cisplatin response in every tissue, has been shown to sensitize glioma cells to these two drugs, while its overexpression reduced their sensitivity [41]. As another example, the knockdown of SALL4 (identified in all tissues) in cancer cell lines has been shown to increase the sensitivity of lung cancer cells [42] and esophageal squamous cell carcinoma cells [43] to cisplatin. Supplementary Table S7 summarizes some of the evidence we curated from literature for the role of different genes identified by TG-LASSO in all tissue types for cisplatin, as an illustration. These examples show the fact that the genes utilized by TG-LASSO in prediction of CDR of patients not only include targets of respective drugs, but also include genes whose expression has been experimentally shown to predict the sensitivity of these drugs: a property necessary for any predictive model of drug response.

Next, we sought to quantify the tissue specificity of identified gene sets for different drugs. For this purpose, we used the Jaccard Distance (JD), which measures the distance of two sets: mutual exclusive sets have JD = 100% and identical sets have JD = 0%. For each drug, we calculated the JD of gene sets identified for that drug in each pair of tissues (Supplementary Table S8). Nine out of 23 drugs had an average JD (calculated across all tissue pairs) of more than 50%. Additionally, there was a high degree of variability in the average JD of different drugs’ gene sets, with Bleomycin having the lowest average JD of 23.4% and Lapatinib having the highest average JD of 65.0% (Supplementary Fig. S3), which suggests a tissue-specific mechanism of action for the latter drug. These results illustrate that TG-LASSO may identify largely different gene sets for a drug from one tissue to another.

### Genes identified for multiple drugs in a tissue are associated with patient survival

We hypothesized that genes that were identified by TG-LASSO as response predictors of many drugs in a single tissue (Supplementary Table S9) may be able to predict the survival of patients who have cancer that originated from that tissue, as they may play a significant role in the development and progress of the disease. To test this, we obtained gene expression values of 4908 primary tumors from 10 different cancer types (corresponding to the tissue types in our study) from TCGA, requiring the data to include at least 170 patients and 20 incidents of deaths for each cancer type (Supplementary Table S10). Then, we clustered the primary tumors of each cancer type into two groups based on the expression of genes identified by TG-LASSO for more than 5 different drugs in the tissue corresponding to that cancer type. We used hierarchical clustering with cosine similarity. Kaplan-Meier survival analysis showed that this clustering approach could separate patients with poor survival from those with better survival (p < 0.05) for 6 out of the 10 cancer types (Fig. 3, Supplementary Fig. S2, Supplementary Table S10). These results provide further evidence in favor of the role of these genes in the progress of the corresponding cancer type.

**Figure 3:**
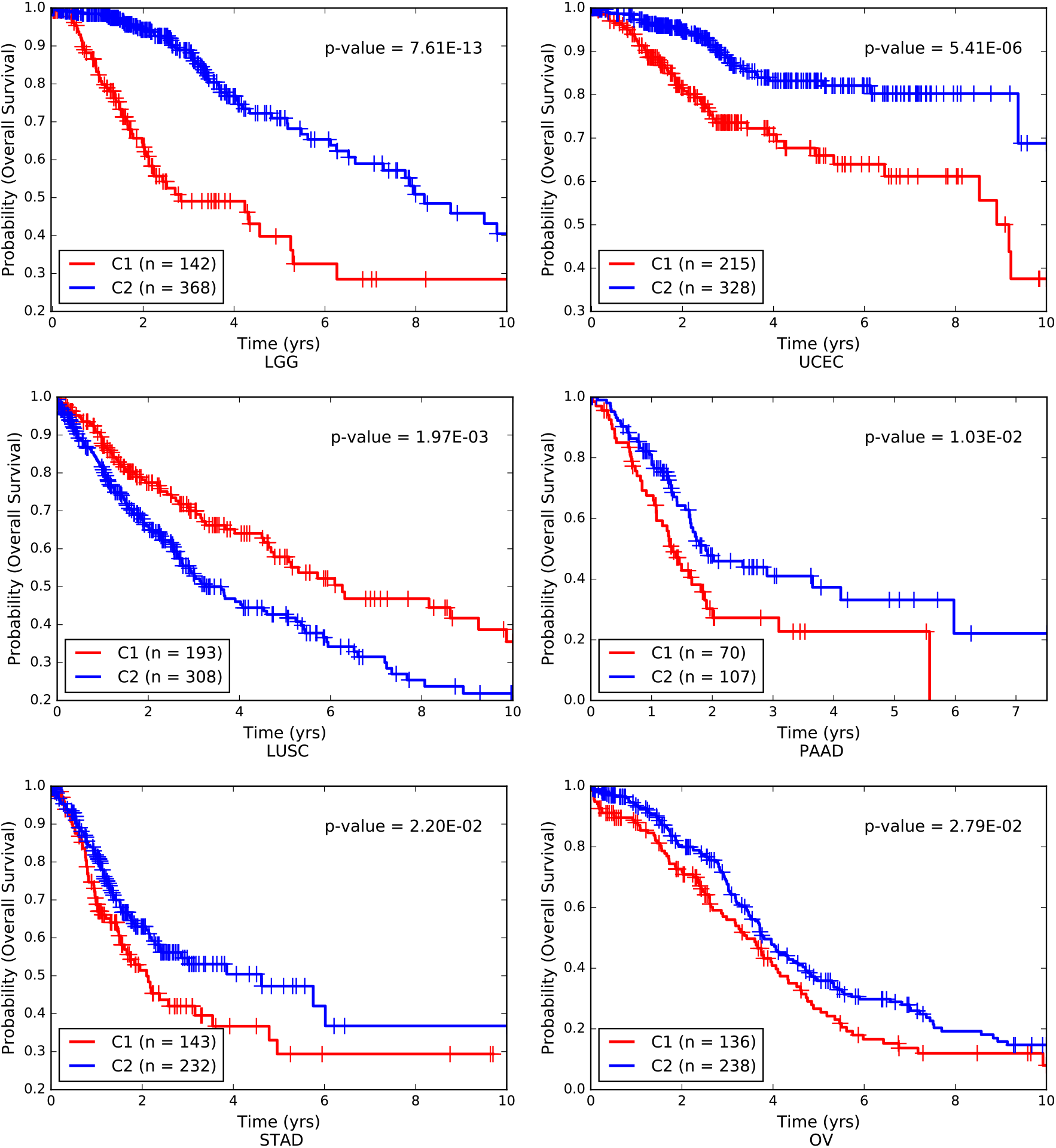
The Kaplan Meier survival analysis results for six cancer types. Patients were clustered based on the expression of genes that were identified by TG-LASSO for more than 5 drugs in the corresponding tissue. The p-value was calculated using a logrank test.

### Functional and pathway enrichment analysis of LGG related genes

Since Kaplan-Meier analysis using Lower Grade Glioma (LGG) patients resulted in the most significant p-value (logrank test, p=7.61E-13), we sought to further characterize the identified genes that resulted in this significant patient stratification using functional and pathway enrichment analysis. For this purpose, we used the KnowEnG’s gene set characterization pipeline [26] and identified 20 gene ontology (GO) terms and two pathways enriched (FDR < 0.05) in this gene set (Supplementary Table S11).

Several of the most significantly enriched GO terms were related to extracellular matrix (ECM), which plays an important role in the infiltration of glioma cells into the brain [44, 45]. Another important GO term was neutrophil degranulation (FDR=2.1E-3). Neutrophils are the most abundant type of white blood cells and the number of infiltrating neutrophils have been shown to be associated with the malignancy of glioma and its drug resistance [46]. In addition, it has been shown that in patients with glioblastoma, neutrophil degranulation is associated with peripheral cellular immunosuppression [47]. Another noteworthy GO term was integrin binding (FDR = 0.037). Integrins are transmembrane proteins that mediate cell adhesion, play an important role in promoting the invasiveness of glioma cells [48], and have been suggested as potential targets with diagnostic and prognostic value in glioma [49]. Several enriched GO terms were related to the activity of endopeptidases and collagen. It has been shown that the level of collagen in glioma patients is increased, and it also plays a key role in promoting the tumor progression [50]. Matrix metalloproteinases (MMPs) are one important class of endopeptidases that are responsible for regulating the turnover of collagens, and their expression and activity has been associated with the progression of human glioma [50, 51]. Finally, ‘response to drug’ was another enriched GO term, which reflects the relevance of the identified genes to the general mechanisms of drug response in a cell.

The enriched pathways included miRNA targets in ECM and membrane receptors (FDR=2.0E-3) and Syndecan-1-mediated signaling (FDR=0.04). Syndecan-1 is a cell surface heparan sulfate proteoglycan and its expression has been shown to be correlated with tumor cell differentiation in various cancers [52]. In addition, its knockdown has been shown to inhibit glioma cell proliferation and invasion and has been suggested as a therapeutic target for glioma [53]. These results support our expectation that the LGG-related gene set not only involves drug response related genes, but also includes those that play important roles in glioma and may act as diagnostic biomarkers or therapeutic targets.

## DISCUSSION

Ideally, a predictive model of CDR should be trained on data obtained directly from patients. Similarly, identification of biomarkers of drug sensitivity has the most potential clinical impact when based on patient data. However, since in practice most patients only receive the “standard of care” treatment based on their specific cancer type, CDR data is scarcely available for the newly approved drugs or drugs that have not yet passed the clinical trial, limiting our ability to decipher the mechanisms of drug sensitivity for these drugs. An alternative approach is to train ML models on preclinical samples (e.g. CCLs) to predict the CDR of patients, then use these predictions to discover novel biomarkers and druggable targets.

Recent large-scale studies that have cataloged the molecular profiles of thousands of CCLs and their response to hundreds of drugs [15–17] are great resources to achieve this goal. In this study, we adopted such an approach and systematically assessed a variety of ML algorithms. Our analyses showed that the CDR of many drugs can be predicted using ML models (especially, regularized linear models) trained on CCLs. However, by evaluating a variety of methods that include auxiliary information (e.g. interaction of the genes, the tissue of origin, etc.), we observed that improving the performance beyond what is achievable using linear models is extremely difficult and requires careful modeling and novel computational techniques. It appears that the way by which auxiliary information is utilized has a large impact: for example, several methods that include the tissue of origin did not improve the results obtained by LASSO, and only TG-LASSO could improve the performance. Additionally, we showed that TG-LASSO identifies tissue-specific gene sets for each drug that include various targets of the drug, genes involved in the drug’s mechanism of action, and genes whose under- or over-expression could sensitize cancer cells to the drug. Moreover, these sets include genes that are involved in cancer progression and are associated with patient survival. These results suggest that in addition to a superior drug response prediction performance, TG-LASSO can identify biomarkers of patient survival and drug sensitivity.

We note that due to the major differences between CCLs and tumors (e.g. the greater heterogeneity of cells in a tumour compared to CCLs), obtaining more accurate results based on classical ML techniques may not be possible. The reason is that classical ML methods assume that the training samples and the test samples are drawn from the same or similar distributions. While batch effect removal and other homogenization and normalization techniques help to alleviate this issue, more realistic preclinical models of cancer are necessary to significantly improve these results. Recent advances in developing human derived xenografts [54] and 3D human organoids [55] may enable developing a more accurate predictive model of CDR in cancer. However, due to the current high cost of these models, a more practical approach is developing computational methods that explicitly model these differences. Such methods must go beyond utilizing bulk gene expression data and take advantage of multi-omics analysis of bulk and single-cell sequencing profiles of samples. Due to the rapid advances in these domains, we expect that large databases of single-cell multi-omics profiles of preclinical and clinical samples and their drug response will become available in the near future.

## METHODS

### Datasets, preprocessing and batch effect removal

We obtained the gene expression profiles (FPKM values) of 531 primary tumor samples of TCGA patients who were administered any of the 23 drugs mentioned earlier. First, we removed genes that contained missing values. We also removed any gene that was not expressed (i.e. FPKM<1) for more than 90% of the samples. Then, we performed a log-transformation and obtained log2(FPKM+0.1) values for each gene. The resulting gene expression matrix contained 19,437 genes and 531 samples. We obtained the CDR of these patients from the supplementary files of [6] (see the original paper for their-approach in curating this data from TCGA). Similarly, we obtained the Robust Multi-array Average (RMA)-normalized basal gene expression profiles and the logarithm of half maximal inhibitory concentration, log(IC50), of 979 cancer cell lines from GDSC (Supplementary Table S1) for 17,737 genes.

To homogenize the gene expression data from these two datasets, we first removed genes not present in both datasets as well as genes with low variability across all the samples (standard deviation < 0.1), resulting in a total of 13,942 shared genes. Then, we used ComBat [20] for batch effect removal to homogenize the gene expression data from TCGA (RNA-seq) and GDSC (microarray). This approach, which has been previously used to successfully homogenize these two data types [21], removed the batch effect present in the gene expression datasets (see Supplementary Fig. S1). For all follow-up analysis, we performed z-score normalization on each gene across all the samples to ensure a mean of zero and a standard deviation equal to one.

For the network-guided analyses, we downloaded four networks of gene interactions in humans from the KnowEnG’s knowledgebase of genomic networks [26] (https://github.com/KnowEnG/KN_Fetcher/blob/master/Contents.md). The details of each network including the number of nodes and edges are provided in Supplementary Table S1.

### Machine learning regression models

The baseline models (Table 1) were all implemented using the Scikit-learn [56] in Python and the hyperparameters were selected using cross validation (using only CCL samples). For the network-based algorithms (Table 4), we used four networks summarized in Supplementary Table S1. We used the normalized graph Laplacian of these networks to run GELnet [31]. This method forces neighboring genes in the graph to have similar weights in order to guide drug response prediction. Specifically, it defines a regularization penalty *R*(*w*) for the standard linear model.

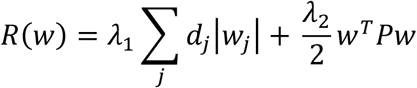

where *d* and *P* are additional penalty weights for individual features and pairs of features, respectively. Our basic GELnet implementation sets *P* = *L* and *d* = 0. Furthermore, we used Network-Induced Classification Kernels (NICK), a method closely related to GELnet. The NICK framework is actually a special case of the GELnet, with *P* = (*I* + *βL*) for some *β* ≥ 0 and *d* = 0. The parameter *β* provides a trade-off between graph-driven regularization and the traditional ridge regression penalty of the SVMs.

In addition to the above methods that utilize the graph Laplacian of each network in the regression algorithm, we used sparse group LASSO (SGL). This method takes a collection of pathways as input and induces sparsity at both the pathway and the gene level to generate the input. We performed community detection on each of the networks in Table 4 by maximizing the modularity using the Louvain heuristics [57] to identify gene sets to be used in the SGL algorithm. We then ran SGL by fitting a regularized generalized linear model with group memberships of genes as deemed by the community detection to predict drug response.

Finally, we developed a heuristic method based on ssGSEA [34] followed by LASSO. In this method, we used ssGSEA to assign a score to each sample for the enrichment of its gene expression profile in communities of each network, obtained earlier. These scores where then used as features to train a LASSO model for prediction of CDR.

### Methods for including tissue of origin in CDR prediction

In the first approach (Method 1 in Table 5), we augmented the gene expression profile of each sample (both CCLs and tumors) with binary features corresponding to different tissues of origin shared between the TCGA and GDSC samples (a total of 13 features). For each sample its tissue of origin was assigned a value of ‘1’, and other tissues were assigned a value of ‘0’. Then, the LASSO algorithm was used to train a drug response model on CCLs and predict the CDR of tumors.

In the second approach (Method 2 in Table 5), we trained different LASSO models for each drug-tissue pair (23 drugs and 13 tissue types). More specifically, to predict the CDR of drug *d* in a tumor of tissue *t*, we trained a LASSO regression model using the IC50 of drug *d* in only cell lines corresponding to tissue *t* (i.e. a subset of the training samples). For tumors originating in tissues without matching training CCLs, we used all the CCLs to train the model.

### Prediction of CDR in cancer tumors using Tissue-guided LASSO

TG-LASSO is a method for predicting the CDR of tumors using the information in *all* training samples (originating from different tissue lineages), while incorporating information on the tissue of origin of the samples. By utilizing all the training samples, it overcomes the lack of generalizability stemming from limited number of CCLs from each tissue type, a major issue in Method 2 above. In addition, by incorporating the information on the tissue of origin of the samples in the training step, it improves the performance of tissue-naïve regression methods, such as those in Table 1.

During training, LASSO minimizes the objective function 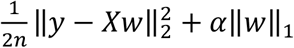, where *n* is the number of training samples, *y* is the response vector of length *n, X* is an *n* × *m* feature matrix (*m* is the number of features), ∥ ∥_2_ denotes the L2 vector norm, ∥ ∥_1_ denotes the L1 vector norm, and *α* is the hyperparameter that determines the sparsity of the model (i.e. number of features used in training). The hyperparameter tuning is usually achieved independent of the structure of the training samples (e.g. their tissue of origin), for example using random cross-validation or a regularization path. However, we and others [58] have shown that including the group structure of data in selecting the hyperparameter is important in assessing the generalizability of regression models. Motivated by these results, even though TG-LASSO utilizes the gene expression and the drug response of all CCLs in training, the hyperparameter *α* is selected in a tissue- and drug-specific manner, as explained below.

Let *D* be the set of all drugs and *T* be the set of all tissues in the test set (i.e. the TCGA dataset). To train a model to predict the CDR of tumor samples from tissue *t* ∈ *T* to drug *d* ∈ *D*, we identify all the training CCLs corresponding to tissue *t* and use them as the validation set. In addition, we use all other CCLs as the training set. Then, the hyperparameter *α* is selected as the one that obtains the best accuracy on predicting the IC50 values of the samples of tissue *t* in the validation set. Designing the hyperparameter-tuning step such that the validation and the test sets have the same tissues of origin ensures that the value of *α* is selected so as to generalize well to the test set. The obtained value of *α* is then used with *all* CCLs (including those from tissue *t*) to fit a model minimizing the LASSO objective function. In the prediction step, this fitted model is then used with the gene expression of tumor samples from tissue *t* to predict their CDR.

### Gene ontology and pathway enrichment analysis

We used the gene set characterization pipeline of KnowEnG analytical platform [26] for this analysis, which utilizes Fisher’s exact test to determine the significance of enrichments. We excluded GOs or pathways with too few genes, focusing only on those with more than 10 members. For the pathway analysis, we used the Enrichr pathways [59] available on KnowEnG. All p-values were corrected for multiple hypothesis testing using Benjamini-Hochberg false discovery rate, available as part of the python module [60].

### Software availability

An implementation of TG-LASSO in python, with appropriate documentation and input files, is available at: https://github.com/emad2/TG-LASSO.

## AUTHOR CONTRIBUTIONS

AE and SS conceived the study and designed the project. AE and EWH designed the algorithms. AE, EWH, AB, and JL implemented the algorithms and performed the statistical analyses of the results. All authors contributed to the drafting of the manuscript and critical discussion of the results. All authors read and approved the final manuscript.

