## Supplemental Methods and Figures for "Tissue-guided LASSO for prediction of clinical drug response using preclinical samples"

### Supplementary Methods, Figures and Tables

#### SUPPLEMENTARY METHODS

##### Within dataset cross-validation performance of network-guided methods compared to LASSO

To test whether network-guided algorithms can improve the within dataset cross-validation performance compared to LASSO, we randomly divided the set of GDSC cell lines into 3 folds of (approximately) equal sizes. One fold was kept aside as the test set and the other two folds were used for training. We used all 23 drugs for our evaluations and focused on NICK as an example. We used Spearman's rank correlation between the predicted IC50 values and the true IC50 values of the test set as the measure of accuracy. Our results showed that in 12 (out of 23) drugs, NICK with the STRING Text Mining and STRING Co-expression networks obtained more accurate results (higher correlation) compared to LASSO. Note that for all three methods, the correlations of predicted and true drug response values were positive and statistically significant ($p<0.05$) for all drugs.

**SUPPLEMENTARY TABLES**

**Table S1:** The summary of data used in this study.

**Table S2:** The detailed drug response prediction performance of baseline methods for each drug.

**Table S3:** The detailed drug response prediction performance of network-based methods for each drug.

**Table S4:** The detailed drug response prediction performance of tissue-based methods for each drug.

**Table S5:** The detailed drug response prediction performance of various methods for drugs whose administration period overlapped with at least another drug (i.e. administered as a combination).

**Table S6:** The list of genes identified by TG-LASSO for each drug, ranked based on the number of tissue in which they were identified.

**Table S7:** The evidence curated from literature for genes identified by TG-LASSO in all tissues for Cisplatin.

**Table S8:** The Jaccard distance between pairs of gene sets corresponding to different tissues identified by TG-LASSO for each drug.

**Table S9:** The list of genes identified by TG-LASSO for each tissue, ranked based on the number of drugs for which they were identified.

**Table S10:** The results of Kaplan Meier survival analysis on TCGA samples using genes identified by TG-LASSO.

**Table S11:** Gene ontology and pathway enrichment results for the top genes identified by TG-LASSO for Brain, used in the survival analysis of LGG samples.

**SUPPLEMENTARY FIGURES**

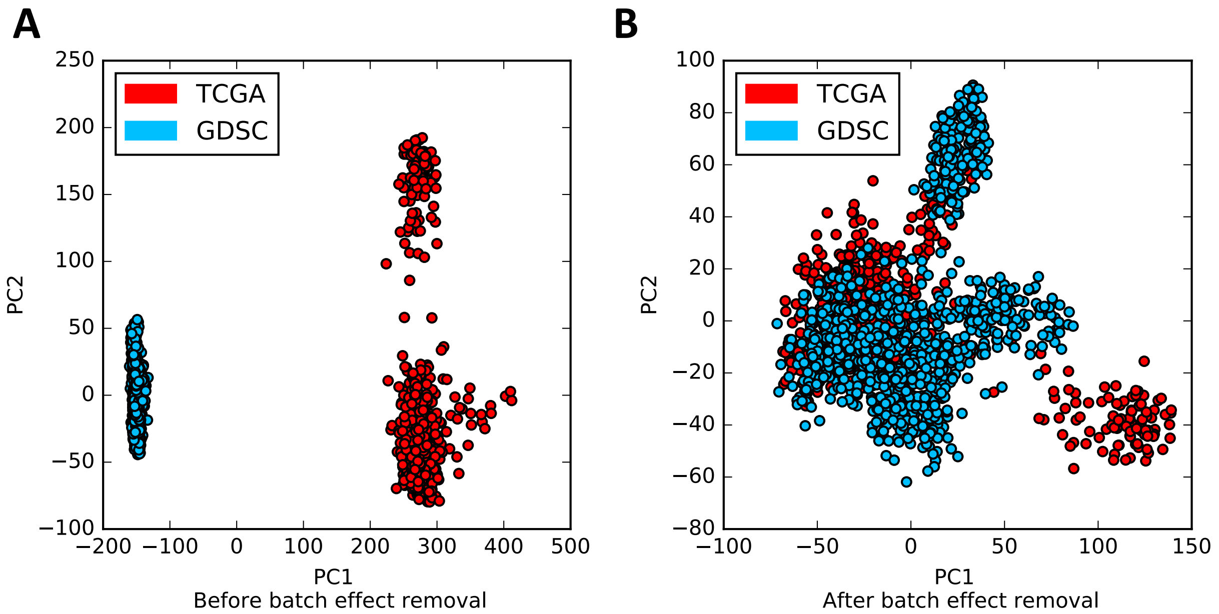


**Figure S1:** The distribution of preclinical as well as tumor samples before (A) and after (B) batch effect removal, depicted using principal component analysis (PCA) of their gene expression.

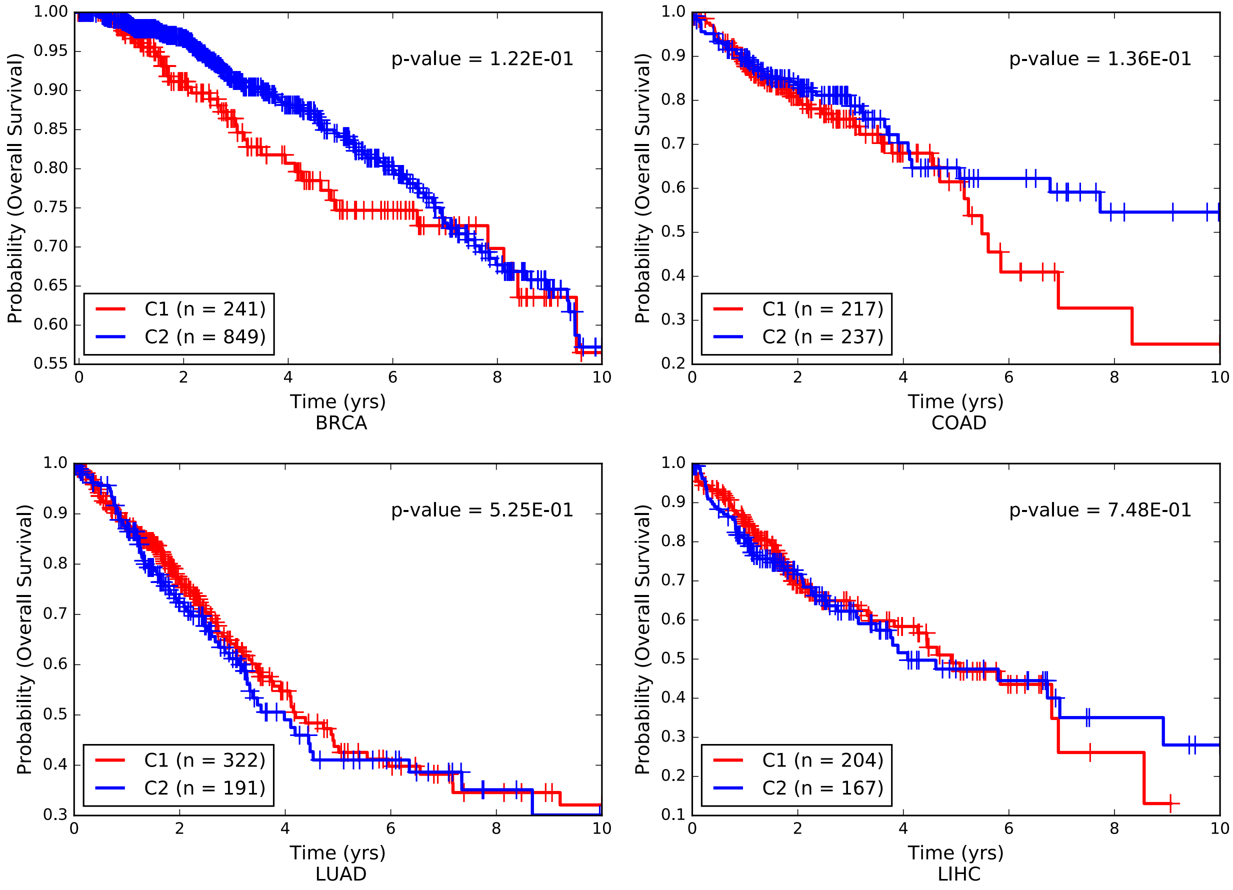


**Figure S2:** The Kaplan Meier survival analysis corresponding to BRCA, COAD, LUAD, and LIHC samples from TCGA. Patients were clustered into two groups using the expression of genes identified by TG-LASSO for more than 5 drugs in each tissue type.

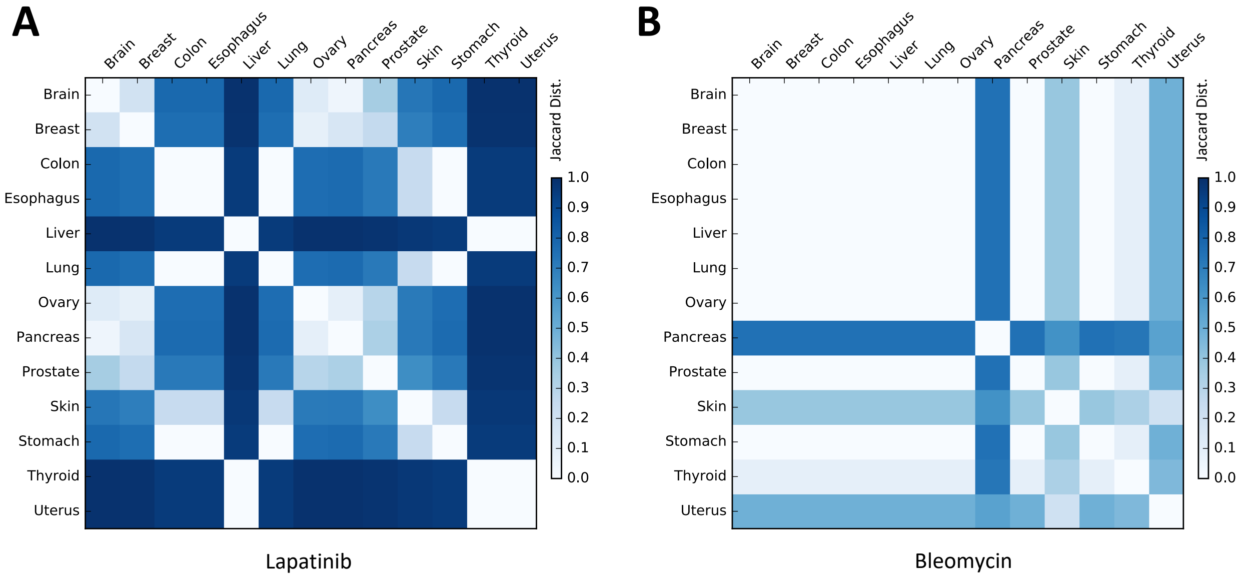


**Figure S3:** The Jaccard distance between pairs of gene sets corresponding to different tissues identified by TG-LASSO for Lapatinib (A), which showed the highest tissue specificity (average Jaccard distance = 65.0%), and Bleomycin (B), which showed the least tissue specificity (average Jaccard distance = 23.4%).
